# Quantitative profiling of 1-Carbon Metabolites, Amino Acids and Precursors, and Plasmalogens in human plasma using Ultra-High-Pressure Liquid Chromatography coupled with Tandem Mass Spectrometry and robotic compound extraction

**DOI:** 10.1101/738864

**Authors:** Stephanie Andraos, Michael Goy, Ben Albert, Martin Kussmann, Eric B. Thorstensen, Justin M. O’Sullivan

**Author notes:** Correspondence: For queries and correspondence, please contact Eric Thorstensen via.

## Abstract

Amino acids (AAs) and one-carbon (1-C) compounds are involved in a range of key metabolic pathways, and mediate numerous health and disease processes in the human body. Previous assays have quantified a limited selection of these compounds. Here, we describe an analytical method for the simultaneous quantification of 37 1-C metabolites, amino acids, precursors, and plasmalogens using reversed-phase ultra-high pressure liquid chromatography coupled with tandem mass spectrometry (UHPLC-MS/MS). Compound extraction from human plasma was tested manually before being robotically automated. The final analytical panel was validated on human plasma samples. Our automated and multiplexed method holds promise for application to large cohort studies.

## 1. Introduction

In the 1930s, Donal Van Slyke and Robert Dillon were some of the first scientists to develop a method for the analysis of free amino acids using crude analytical methods (1). Since then, the comprehensive analysis of low-molecular weight metabolites (*i.e*. metabolomics) has progressed thanks to advanced technology, in particular the coupling of chromatography and mass spectrometry, to enable their accurate quantitation in biological samples (2,3). Several metabolomics methods have been developed for amino acids and/or 1-C metabolism compound quantification in different body fluids (*e.g*. plasma, cerebrospinal fluid, red blood cells) (4,5). However, on their own, each of these methods only quantifies a limited selection of metabolites (4,5).

### 1.1 Amino acids and 1-C metabolites in human health

Amino acids and 1-C metabolites are central to physiological processes in the human body (*e.g*. nutritional metabolism, molecular, endocrine and neurological functions) (6). Due to their ubiquitous involvement in biological processes, quantifying the concentration of amino acids and 1-C compounds in human plasma can inform on the physiological status of an individual. Specific metabolic profiles have been linked to health and disease outcomes (7,8). Estimating nutritional status using dietary reporting alone is unreliable (9). Therefore, an objective quantitation of nutritional metabolites is of utmost importance, to strengthen our current understanding of human physiology and nutritional metabolism.

### 1.2 The use of human plasma for quantitative metabolomics

Plasma is the non-cellular component of human blood. As such, plasma contains virtually all human proteins (*e.g*. blood clotting, binding proteins) representing those expressed throughout the tissues in addition to glucose, vitamins, and other nutrients. Blood collection from human subjects is relatively easy and non-invasive making it particularly appealing in both clinical and research settings (10). In a healthy population, blood metabolites (*e.g*. amino acids) typically maintain homeostatic levels when in a fasted state, but may fluctuate substantially in the post-prandial state (11). Therefore, physiological extrapolations need to be made cautiously when interpreting the results of plasma metabolomics studies, as these reflect either states of homeostasis (in the fasted state), a postprandial response (in the fed state), or metabolic exchanges between tissues. Notably, plasma-tissue correlations of metabolites are not always reliable (11). For example, metabolites quantified in plasma are typically not able to be assigned to cells or tissues of origin. However, an increasing level of biological understanding of tissue-specific pathways has allowed for the development of mathematical models, extrapolating organ-specific metabolite levels based on their plasma measurements (12,13).

Here we present an automated method that combines a robotic extraction of plasma samples with reversed-phase ultra-high-pressure liquid chromatography coupled with tandem mass spectrometry (UHPLC-MS/MS) to quantitate 37 1-C metabolites, amino acids and precursors, as well as plasmalogens in a single plasma sample. This method has wide applicability for metabolomic studies using non-invasive sample collection methods.

## Materials and Methods

### Ethics

Plasma samples used for method validation were obtained from the Fish Oil in Pregnancy trial, approved by the Northern A ethics Health & Disabilities Committees (Trial number 17/NTA/154). Participants provided written informed consent, and the trial was conducted according to the principles of the declaration of Helsinki.

### Materials

Materials used throughout the experiment are detailed in Supplementary table 1.

### Plasma sample collection

Whole blood was collected in lithium heparin and EDTA blood collection tubes. Plasma was separated by centrifugation (1300 x g, 4^⍰^C, 15 min), then stored (– 80^⍰^C).

### Standard preparation and quality controls

A labelled amino-acid premix solution containing: L-Arginine:HCL (^13^C6, 99%; ^15^N4, 99%); L-Aspartic acid (^13^C4, 99%; ^15^N, 99%); L-Cystine (^13^C6, 99%; ^15^N2, 99%); L-Glutamic acid (^13^C5, 99%; ^15^N, 99%); L-Histidine:HCL:H2O (<5% D) (U-^13^C6, 97-99%; U-^15^N3, 97-99%); L-Isoleucine (^13^C6, 99%; ^15^N, 99%); L-Leucine(^13^C6, 99%; ^15^N, 99%); L-Methionine (^13^C5, 99%; ^15^N, 99%); L-Phenylalanine (^13^C9, 99%; ^15^N, 99%); L-Proline (^13^C5, 99%; ^15^N, 99%); L-Threonine (^13^C4, 97-99%; N, 97-99%); L-Tyrosine (^13^C9, 99%; 15N, 99%); and L-Valine (^13^C5, 99%; N, 99%) (*Cambridge Isotope*) was diluted to 33 µM in 1% ascorbic acid MilliQ® H_2_O (Millipore). Stock solutions of labelled 1-C compounds were prepared from their corresponding powders of: Homocysteine d4 (Novachem (CIL)); Cystathionine d4 (Novachem (CIL)); S-Adenosylhomocysteine d4 (Sapphire bioscience (Cayman)); 5-Methyltetrahydrofolate d3 (Sapphire Bioscience Pty Ltd); Betaine d3 (SciVac PTY. Ltd. (CDN)), Dimethylglycine d3 (SciVac PTY. Ltd. (CDN)); and Taurine d4 (SciVac PTY. Ltd. (CDN)) diluted in 0.1 M HCl for all standards, except for 5-MTHF d3, which was diluted in 10 mM ammonium acetate (NH_4_OAc) with 10% ascorbic acid and 2% TCEP, at concentrations ranging from 0.5 to 5 mM. These stock solutions were then diluted to 33 µM in 1% ascorbic acid MilliQ® H_2_O. A 1:1 mixture of labelled AAs and 1-C compounds was used as the internal standard solution.

Dilution series’ of unlabelled amino acids (Kit No. LAA-21, SIGMA Chemical company, USA) and 1-C metabolites were prepared at concentrations ranging from 1µM to 1000µM (Supplementary tables 2 and 3), and mixed in a 1:1 ratio to create standard 8 (S8; Table 1). The unlabelled standards consisted of Lysine, Taurine, OH-Proline, Proline, Aspartic acid, Glutamine, Glycine, Serine, Asparagine, Glutamic acid, Dimethylglycine, Citrulline, Proline, Betaine, Threonine, Cysteine, Ethanolamine, Alanine, Aminoadipic acid, Homocysteine, Ornithine, Valine, Carnitine, Tyrosine, Choline, Methionine, Betaine, Cystathionine, Phenylalanine, Isoleucine, Leucine, Histidine, Arginine, S-Adenosylhomocysteine, Tryptophan, S-Adenosylmethionine, 1-Methylhistidine, 3-Methylhistidine, and 5-Methyltetrahydrofolate (all purchased from Sigma Aldrich). Dilutions of unlabelled mixes were prepared to create a wide dynamic range of concentrations that mirrors known physiological plasma concentrations (Table 1).

**Table 1:** Calibration curve concentrations

| Table 1: Calibration curve concentrations |  |  |  |  |  |  |  |  |  |
| --- | --- | --- | --- | --- | --- | --- | --- | --- | --- |
| Compound | S1 | S2 | S3 | S4 | S5 | S6 | S7 | S8 | Unit |
| Homocysteine | 0.2 | 1 | 2 | 5 | 10 | 20 | 40 | 100 | µM |
| Choline | 0.4 | 2 | 4 | 10 | 20 | 40 | 80 | 200 | µM |
| Dimethyl glycine |  |  |  |  |  |  |  |  | µM |
| Glutamic acid |  |  |  |  |  |  |  |  | µM |
| Ornithine |  |  |  |  |  |  |  |  | µM |
| Methionine |  |  |  |  |  |  |  |  | µM |
| Tyrosine |  |  |  |  |  |  |  |  | µM |
| Isoleucine |  |  |  |  |  |  |  |  | µM |
| Phenylalanine |  |  |  |  |  |  |  |  | µM |
| Tryptophan |  |  |  |  |  |  |  |  | μM |
| Ethanolamine |  |  |  |  |  |  |  |  | μM |
| Amino adipic acid |  |  |  |  |  |  |  |  | μM |
| Aspartic acid |  |  |  |  |  |  |  |  | μM |
| Citrulline |  |  |  |  |  |  |  |  | μM |
| 1-methylhistidine |  |  |  |  |  |  |  |  | μM |
| 3-methylhistidine |  |  |  |  |  |  |  |  | μM |
| OH-Proline |  |  |  |  |  |  |  |  | μM |
| Histidine |  |  |  |  |  |  |  |  | μM |
| Asparagine |  |  |  |  |  |  |  |  | μM |
| Cysteine | 0.5 | 2.5 | 5 | 12.5 | 25 | 50 | 100 | 250 | μM |
| Betaine |  |  |  |  |  |  |  |  | μM |
| Leucine | 0.6 | 3 | 6 | 15 | 30 | 60 | 120 | 300 | μM |
| Cystathionine | 1 | 5 | 10 | 25 | 50 | 100 | 200 | 500 | nM |
| S-Adenosylhomocysteine (SAH) |  |  |  |  |  |  |  |  | nM |
| S Adenosyl-L-methionine (SAM) |  |  |  |  |  |  |  |  | nM |
| Taurine |  |  |  |  |  |  |  |  | μM |
| Alanine |  |  |  |  |  |  |  |  | μM |
| Serine |  |  |  |  |  |  |  |  | μM |
| Glycine |  |  |  |  |  |  |  |  | μM |
| Glutamine |  |  |  |  |  |  |  |  | μM |
| Proline |  |  |  |  |  |  |  |  | μM |
| Lysine |  |  |  |  |  |  |  |  | μM |
| Arginine |  |  |  |  |  |  |  |  | μM |
| Valine |  |  |  |  |  |  |  |  | μM |
| Threonine |  |  |  |  |  |  |  |  | μM |
| Carnitine |  |  |  |  |  |  |  |  | μM |
| 5-Methyltetrahydrofolic acid (5-MTHF) | 5 | 25 | 50 | 125 | 250 | 500 | 1000 | 2500 | nM |

Stripped human plasma (SeraCare Life Sciences Inc.) was used as a quality control (QC) matrix to calculate compound recoveries. Triplicate QCs of stripped plasma at baseline and stripped plasma spiked with 70 µL of calibration standards (S7 and S8) were used for quality control checking.

### Automated solid-phase extraction (SPE)

An Eppendorf robot fitted with both a thermal mixer and vacuum manifold (EpMotion 5075vt, Germany) was programmed using the epBlue Client v. 40.6.2.6 software to automate compound extractions from plasma samples (Figure 1). The protocol was developed to extract one 96-well plate of samples including standards, blanks and QCs. Internal standards, quality controls, calibration standards, the protein precipitation solution, the TCEP solution, 1% ascorbic acid MilliQ® H_2_O, and plasma samples were all placed in the robot.

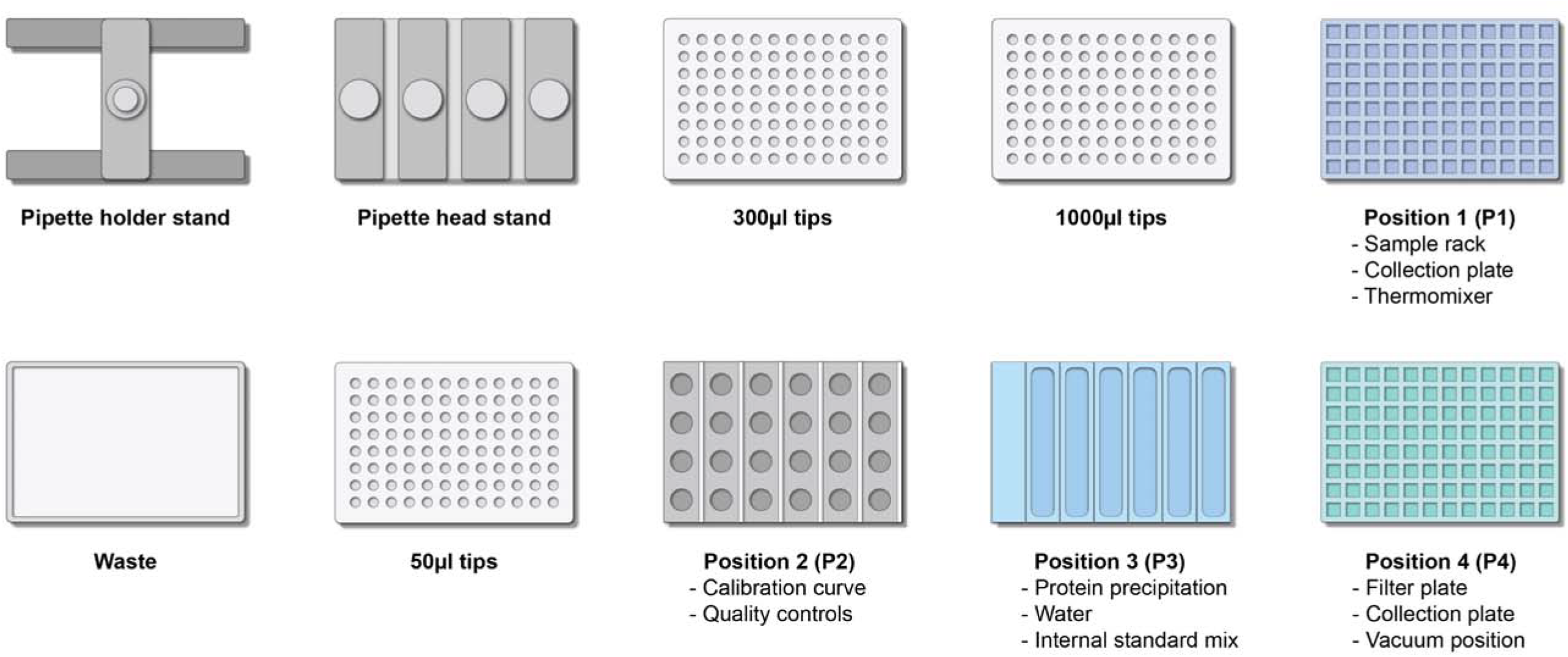

### Protein precipitation

Firstly, 300 µL of 1% (v/v) formic acid in LC-Grade MeOH (in P3) is transferred into the all wells (in P4) of the 96-well IMPACT® protein precipitation plate (Phenomenex®) fitted onto a square 96-well (2ml) collection plate (Phenomenex®). A volume of 100 µL of 1% ascorbic acid in MilliQ® H_2_O is dispensed into the first two wells as blanks with and without internal standards. The internal standard solution (20 µL from P3) is dispensed into the remaining wells of the plate. Standards (100 µL; in P2) and plasma samples (100 µL; in P1) are transferred into the corresponding wells of the plate. Quality controls (QCs; 100 µL in triplicate) are dispensed from P2 into three different locations across the plate, to enable quantification of QC fluctuations at different injection times across the 96-well plate.

### Mixing and vacuum extraction

Once sample transfer is completed, the rack containing plasma samples in P1 is removed from the robot, recapped, and stored (−80°C). The protein precipitation filtration plate and catch plate are then placed on a thermomixer and agitated (5 min, 800rpm, room temperature). Once mixed, the plate is placed on the vacuum stand. A vacuum (450mbar, 10 min) is applied and the extracted compounds are collected in the catch plate.

### Addition of the reducing agent TCEP

After precipitation of the extracted compounds, the filter plate is discarded and 100 µL of Tris (2-carboxyethyl) phosphine (TCEP) is transferred from P3 to all wells of the collection plate. TCEP reduces the disulphide bonds within cystine and homocystine, so they can be quantified as cysteine and homocysteine.

### Mixing and final dilution

The collection plate is placed on P1 for mixing (15 min, 800 rpm, room temperature), to provide sufficient contact time between the TCEP solution and the extracted compounds of interest for optimal compound reduction. A volume (200 µL) of 1% ascorbic acid in MilliQ® H_2_O is then added to each well. The plate is capped using a 96 square sealing mat (Phenomenex®), mixed (5 min, 800rpm, room temperature), and placed in the UHPLC system auto-sampler (10°C) for analysis.

### UHPLC/MS-MS working conditions

Ultra-high-pressure liquid chromatography coupled with tandem mass spectrometry was performed using a Vanquish UHPLC system, coupled with a TSQ Quantiva triple quadrupole mass spectrometer (Thermo Scientific), using a heated electrospray ionisation source (H-ESI) in positive ionization mode. A Kinetex® EVO C18 100 Å 150×2.1mm 1.7µm column (Phenomenex®) at 40°C, coupled with a Krudkatcher (Phenomenex®) pre-column filter, was used for chromatography. A flow of 400 µL/min starting at 2% Acetonitrile and 98% mobile phase consisting of 5mM perfluorohexanoic acid (PFHA) in MilliQ® H_2_O was applied to the column, compounds of interest were eluted using an increasing acetonitrile gradient (Table 2). The sample injection volume was 7 µL, and the run time was 15.5 minutes.

**Table 2:** LC flow gradient

| Table 2: LC flow gradient |  |  |
| --- | --- | --- |
| Time | Flow (ml/min) | Percentage Acetonitrile (%) |
| 0 | 0.4 | 2 |
| 0.5 | 0.4 | 2 |
| 2.5 | 0.4 | 15 |
| 4.0 | 0.4 | 15 |
| 6.3 | 0.4 | 20 |
| 7.3 | 0.4 | 20 |
| 9.5 | 0.4 | 95 |
| 11.5 | 0.4 | 95 |
| 11.6 | 0.4 | 2 |
| 15.5 | 0.4 | 2 |

### Data processing

Data processing was carried out using Xcalibur version 4.0.27.19 (Thermo). Compounds were assigned specific retention times (based on the injection of pure compounds), and labelled standards for quantitation purposes. Unlabelled compounds that did not have labelled counterparts were assigned other labelled compounds with similar properties (*e.g*. SAM and SAH were both assigned SAH d4 as an internal standard for quantification). The range of expected concentrations of the compounds in standards S1-S8 (Table 1) were also included.

### Plasma sample validation

The process was validated using stripped plasma and pooled plasma samples from pregnant women taking part in the Fish Oil in Pregnancy study. The validation included: 1) measures of specificity and selectivity based on retention times; chromatographic separations and spectral detection; 2) measures of matrix effects and standard recoveries by comparing pure stripped plasma versus spiked stripped plasma samples; and 3) measures of reproducibility by calculating coefficients of variation for multiple injections of the independent extractions of the same pooled plasma sample within and across multiple assay runs.

## Method development

### Spectral Tuning

Compounds were tuned on a TSQ Quantiva mass spectrometer using the TSQ Quantiva Tune Application v. 2.0.1292.15 software. Compound tuning identifies the lens’ radio frequencies (RF) and respective collision energies required to focus and fragment a parent ion into its daughter ion(s). Each unlabelled standard was diluted from the stock solution in 1% ascorbic acid of MilliQ® H_2_O. The tuning solution of each individual compound was loaded into a tuning syringe. A LC flow (100-200 µl/min) was introduced into the mass spectrometer via ‘T’ connector to mimic standard working conditions. After each standard was infused into the mass spectrometer via a syringe, a stepwise entry of ascending collision energies was applied to identify the ideal conditions for the detection of the most abundant daughter ion of the corresponding parent ion of interest. The mass of the most abundant daughter ion was identified along with its RF lens voltage and collision energy for each compound.

### UHPLC/MS-MS Instrument setup

The instrument setup consisted of two stages: 1) chromatographic parameters; and 2) spectral conditions. Chromatographic flow gradient parameters were optimised to maximise compound separation whilst minimising assay run time. The optimised flow gradient is detailed in Table 2. Spectral parameters were optimised through an instrument setup that was drafted and edited throughout the method development process, including all compound names, their corresponding collision energies, RF lens, parent and daughter mass transitions, as well as expected retention times and windows.

## Chromatographic separations

### Chromatographic columns

Solutions of each compound and internal standard were tested to determine their retention times under the gradient used. Initial work used a Kinetex® 2.6µm F5 100 Å 150×2.1 mm column (Phenomenex^®^) column to test chromatographic separations. However, some compound peaks showed poor resolution. A smaller particle size column, the Kinetex® 1.7µm EVO C18 100 Å 150 x2.1 mm column (Phenomenex^®^) showed improved resolution, and was used for chromatographic separations instead.

### Ion-pairing reagents

Ion-pairing reagents are mobile phase additives pairing with highly ionic compounds to improve their adsorption to the stationary phase of the chromatographic column. Heptafluorobutyric acid (HFBA) was initially tested as an ion-pairing reagent, but exhibited poor retention of some compounds in this method. Perfluorohexanoic acid (PFHA) improved compound retention on the chromatographic column (5mM of PFHA in MilliQ® H_2_O) and was subsequently used instead.

## Results and discussion

Parent-product ion transitions, collision energies, and RF lens of all compounds were obtained (Table 3). The compounds included in this panel are present in a wide range of metabolic networks and have been associated with health and disease outcomes (*i.e*. the 1-carbon metabolism, the urea cycle, ketogenic and glucogenic amino acids involved in the tricarboxylic acid (TCA) cycle, branched-chain amino acid (BCAA) metabolism, Trimethylamine N-Oxide (TMAO) metabolism, and gamma-aminobutyric acid (GABA) metabolism (14–21)). It has been argued that quantifying single blood biomarkers that are robustly associated with an outcome of interest (*e.g*. hsCRP, HbA1c) may be more clinically relevant for bigger picture recommendations and the development of public health policies (22). Whilst robust biomarkers must be profiled for diagnostic purposes, linking nutritional metabolites (*e.g*. 1-C metabolites, and amino acid profiles), to pre-established markers of health outcomes remains crucial for the characterisation of potentially targetable - underlying mechanistic layers linking exposure to outcome (19,23). Of these mechanisms, epigenetic processes (such as DNA and histone methylation) are largely dependent on the availability of methyl (-CH_3_) groups released by methyl donors produced from the one-carbon metabolism (24–28). Briefly, this analytical panel measures metabolites that are at the intersection of lifestyle, nutrition, metabolism, and epigenetics (24–28). Whilst some of these compounds are unstable, prone to oxidation, and tend to degrade in samples stored for long periods, the exhaustiveness of our panel allows for a biologically relevant, yet still practically applicable coverage of multiple biological networks in a single 100µl plasma sample.

**Table 3:**
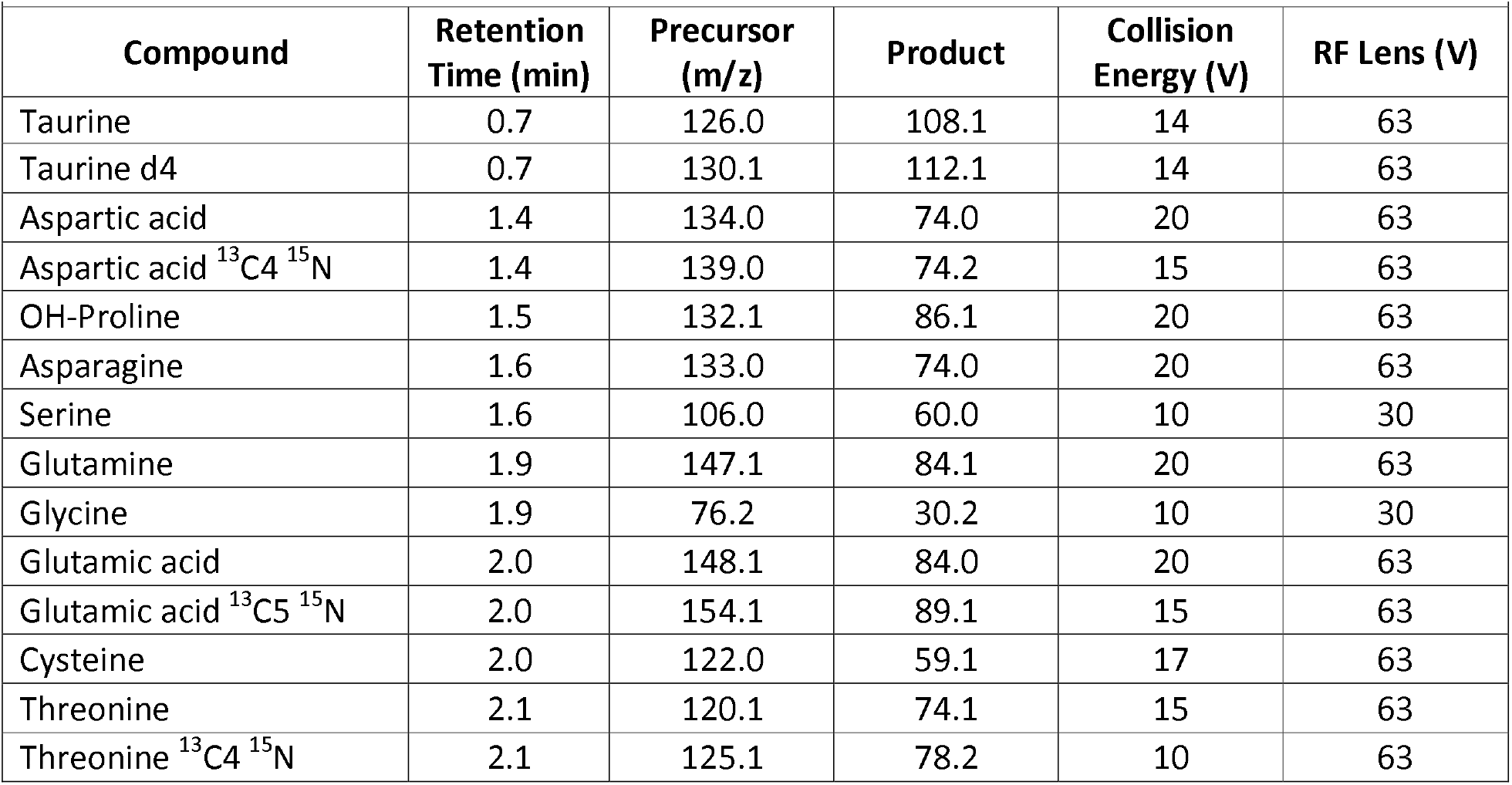
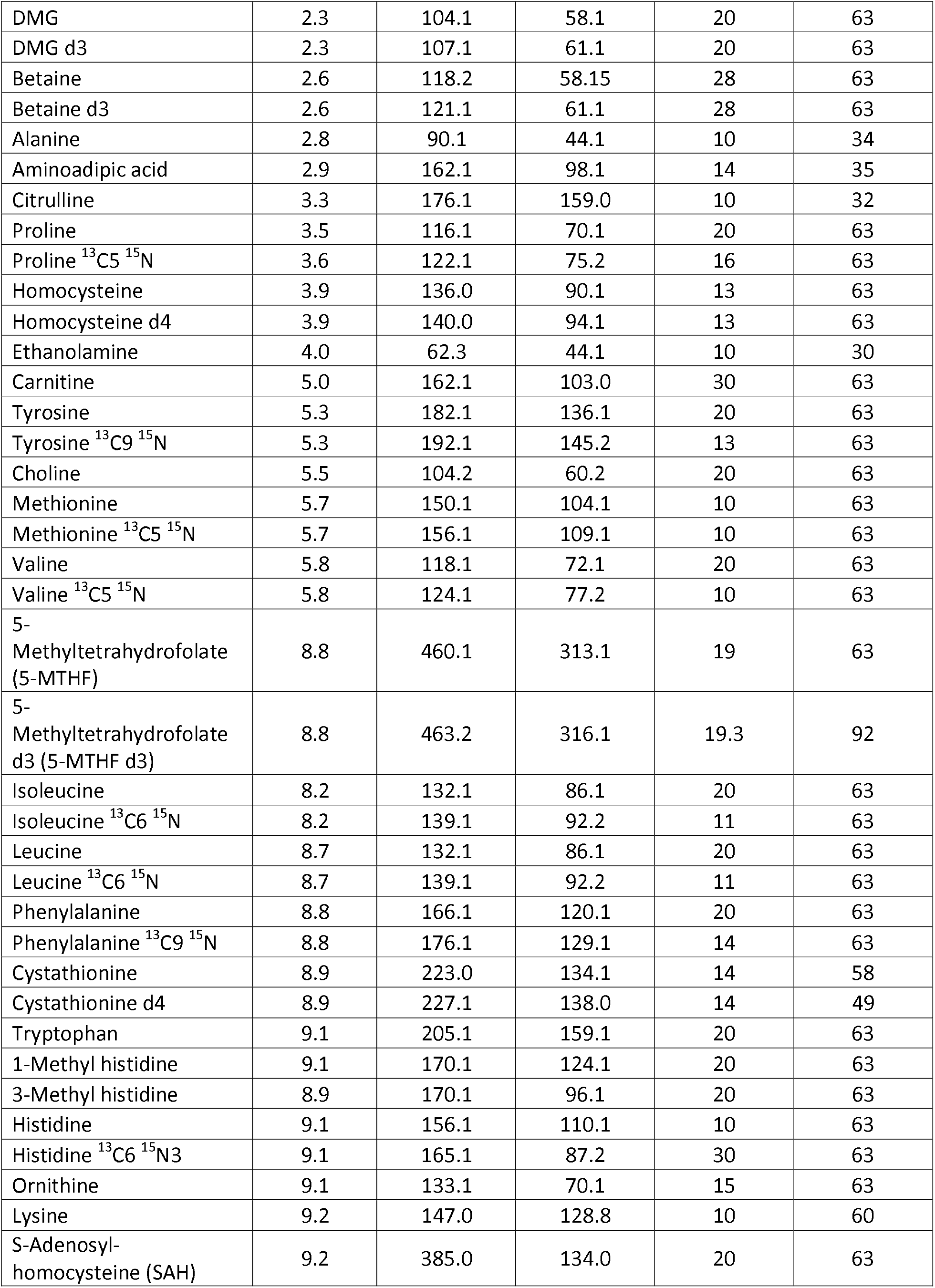
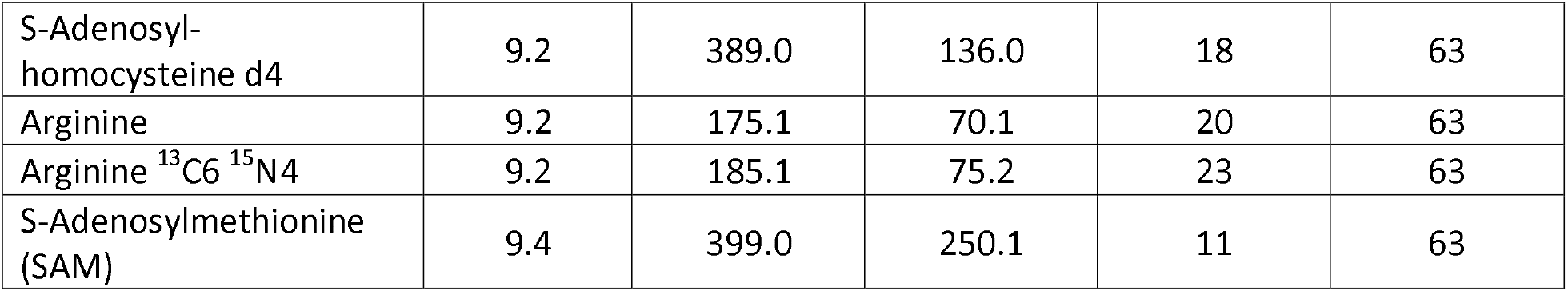
Single-reaction monitoring (SRM) transitions and compound retention times

### Compound recoveries and inter-assay coefficients of variations

A number of plates were tested during method development. Quality control (QC) results of 96-well plates profiled by the same user between March and July 2019 were reported in the following. QCs consisted of: QC-1 (stripped plasma at baseline), QC-2 (stripped plasma spiked with S7 concentration), QC-3 (stripped plasma spiked with S8 concentration), and QC-4 (real pooled plasma samples). Triplicate QCs were injected at three different locations of a 96-well plate to track compound performance and stability across a plate taking 26-30h to run. The injection time separating each set of QCs was approximately 10 hours allowing for a good representation of real changes occurring in samples throughout the plate. Compound recoveries reported were calculated by comparing baseline stripped plasma with spiked stripped plasma (Table 4). Given that each compound behaves slightly differently across plates profiled months apart, outliers were excluded from the total recovery calculations (1, 2, and 3). The exclusion criteria were ± 2sds for most compounds, and ± 1sd for those that tended to fluctuate more markedly over 5 months of runs (Asparagine, Cysteine, Betaine, Ethanolamine, and S-Adenosylhomocysteine). The reported recoveries were mostly for the S7 spike, given that it is closer to physiological ranges of compound concentrations in plasma, and is therefore more biologically relevant. For metabolites exhibiting lower sensitivity or higher baseline levels in stripped plasma, a higher spike concentration S8 was used for this calculation. These compounds were Cysteine, Betaine, Alanine, S-Adenosylhomocysteine, Asparagine, and Glutamic Acid. All compounds recovered within two standard deviations of 100% recovery, highlighting a reliable quantitation of compounds of interest (Table 4). 5-Methyltetrahydrofolate (5-MTHF) had an acceptable recovery but results for this compound were inconsistent across runs. This is a clear downfall of a complex panel attempting to quantify multiple compounds simultaneously. Regardless of the acceptable recoveries in some plates, the inconsistencies between plates highlight the need to quantify folate derivatives such as 5-MTHF in a specialized method with minimal extraction time and stringent, light-protected blood collection methods given the highly unstable and oxidative nature of these compounds (29,30).

**Table 4:** Compound recoveries and reproducibility in stripped and pooled plasma

| **Compound** | **Recoveries in stripped plasma (Rounded %)** | **Mean levels in pooled fasted plasma** | **Unit** | **Coefficient of variation of pooled plasma concentration within a plate or intra-assay variability (Rounded %)** | **Coefficient of variation of pooled plasma concentration across plates or inter-assay variability (Rounded %)** |
| --- | --- | --- | --- | --- | --- |
| Lysine | 105 | 178 | μM | 11 | 19 |
| Taurine | 111 | 33 | μM | 14 | 25 |
| OH-Proline | 91 | 8 | μM | 6 | 12 |
| Aspartic acid | 96 | 4 | μM | 7 | 13 |
| Glutamine | 102 | 678 | μM | 6 | 22 |
| Serine | 90 | 79 | μM | 9 | 8 |
| Glycine | 102 | 144 | μM | 10 | 13 |
| Asparagine | 87 | 31 | μM | 5 | 8 |
| Glutamic acid | 91 | 40 | μM | 4 | 7 |
| DMG | 92 | 2 | μM | 5 | 20 |
| Citrulline | 90 | 15 | μM | 10 | 20 |
| Proline | 91 | 141 | μM | 4 | 6 |
| Betaine | 79 | 13 | μM | 7 | 13 |
| Threonine | 84 | 138 | μM | 5 | 9 |
| Cysteine | 77 | 47 | μM | 10 | 17 |
| Alanine | 91 | 260 | μM | 7 | 10 |
| Homocysteine | 77 | 1 | μM | 14 | 20 |
| Ornithine | 83 | 14 | μM | 22 | 13 |
| Valine | 92 | 157 | μM | 4 | 8 |
| Carnitine | 109 | 25 | μM | 11 | 22 |
| Tyrosine | 89 | 36 | μM | 4 | 5 |
| Choline | 116 | 9 | μM | 10 | 15 |
| Methionine | 95 | 19 | μM | 4 | 7 |
| Cystathionine | 97 | 99 | nM | 14 | 18 |
| Phenylalanine | 92 | 48 | μM | 3 | 2 |
| Isoleucine | 91 | 43 | μM | 4 | 6 |
| Leucine | 90 | 80 | μM | 4 | 8 |
| Histidine | 100 | 76 | μM | 7 | 9 |
| Arginine | 96 | 60 | μM | 5 | 7 |
| S-Adenosylhomocysteine (SAH) | 79 | 31 | nM | 21 | 15 |
| S-Adenosylmethionine (SAM) | 99 | 61 | nM | 14 | 18 |
| Tryptophan | 95 | 44 | μM | 12 | 22 |
| 1-Methyl Histidine | 81 | 3 | μM | 6 | 19 |
| 3-Methyl Histidine | 88 | 5 | μM | 8 | 9 |
| Aminoadipic acid | 81 | 3 | μM | 9 | 10 |
| Ethanolamine | 128 | <LOQ* | μM | N/A | N/A |
| 5-Methyltetrahydrofolate (5-MTHF) | 91 | 27 | nM | N/A | N/A |

### Mean concentration calculations across assays

Compound concentrations were reported as average concentrations of pooled plasma QCs within the ±2 sds of the average across plates profiled between March and July 2019. Compounds reported as below LOQ (limit of quantification) were those exhibiting concentrations below the lowest calibration curve standard (S1).

### Reproducibility, inter- and intra-assay coefficients of variation

Compound reproducibilities were calculated by comparing compound concentrations 1) at different locations on the same 96-well plate to monitor fluctuations across a run (every 10h); and 2) between plates by calculating standard deviations of pooled plasma QCs (QC4) injected in different plates, across several months. Inter-and intra-assay variabilities were reported (Table 4), most of which were below our 20% cut-off, pinpointing a stable and consistent quantitation of these metabolites for the same samples both within and between assays. For compounds exhibiting low concentrations in pooled plasma, inter- and intra-assay reproducibility was reported for spiked stripped plasma samples exhibiting higher concentrations.These include: Aspartic acid, OH-Proline, Aminoadipic Acid, and 3-Methylhistidine. The rationale behind that substitution is that very low concentrations are bound to give falsely higher coefficients of variations, therefore falsely lower reproducibility.

## Conclusion

We have developed, robotically automated, and validated a methodology for the simultaneous quantitation of an extensive metabolic panel of 1-C compounds, amino acids and precursors, as well as plasmalogens in human plasma. Our method has two key advantages over existing methods: Firstly, our technique is highly reproducible and robust due to the robotic automation of sample preparation and compound extraction. The ability to measure metabolites such as these directly from their sample vial with no manual ‘intervention’ is attractive from an analysis and laboratory staffing point of view. Secondly, our method has a greater coverage of compounds from a number of metabolic pathways, when compared to previous methods (4,5), which provides simultaneous information on interconnected biological processes rather than focusing on just one. These advantages mean that this method has the potential for wide-spread application to large cohorts in a wide range of fields (7,31–33).

## Acknowledgments

This study was funded by a Faculty Research Development fund grant to MK, and a MBIE Catalyst grant (The New Zealand-Australia LifeCourse Collaboration on Genes, Environment, Nutrition and Obesity (GENO); UOAX1611; to JOS). SA is the recipient of a New Zealand International Doctoral Research Scholarship 2017. Samples used in this study were collected as part of the Fish Oil in Pregnancy Study, which was funded by a Health Research Council Emerging Researcher First Grant and by a project grant from the A Better Start National Science Challenge and Cure Kids.

The authors would like to thank PhD student Vidit Satokar for collecting the samples as part of the Fish Oil in Pregnancy study, and the participants of the trial.

