## Supplementary material for "Quantitative profiling of 1-Carbon Metabolites, Amino Acids and Precursors, and Plasmalogens in human plasma using Ultra-High-Pressure Liquid Chromatography coupled with Tandem Mass Spectrometry and robotic compound extraction"

| **Table 1: List of materials and corresponding suppliers** | | |
| --- | --- | --- |
| **Chemical** | **Supplier** | **Catalogue number** |
| LC-grade Acetonitrile | LiChrosolv (Germany) | 1.00030.2500 |
| LC-grade Methanol | EMSURE (Germany) | 1.06007.2500 |
| Perfluoroheptanoic acid (PFHA) | Sigma Aldrich | 342041 |
| Ascorbic acid | Scharlau Chemie (Barcelona). | AC0515 |
| Impact protein precipitation 96-well plate | Phenomenex® | CEO-7565 |
| 96-well collection plate (square 2ml/well) | Phenomenex® | AH0-7193 |
| Eppendorf tips (50µL, 300µL and 1000µL) | Eppendorf | EP0030014421, EP0030014464, EPP0030003993 |
| Amino acid labelled standard mix | Cambridge isotope laboratories, Inc. | MSK-A2-1.2 |
| DL-Homocysteine-d4 | Cambridge Isotopes | DLM-8259-0.1 |
| Cystine d4 | Cambridge Isotopes | DLM-9812-0.5 |
| DL Homocysteic acid | CDN Isotopes | D-5646 |
| D Cystathionine | CDN Isotopes | D-7717 |
| Betaine d3 | CDN Isotopes | D-6303 |
| DimethylGlycine d3 | CDN Isotopes | D-7024 |
| L-Cysteine | Sigma Aldrich | C 7755 |
| Homocysteine | Sigma Aldrich | 69453-10MG |
| Cystathionine | Sigma Aldrich | C7505 |
| S-Adenosylhomocysteine | Sigma Aldrich | A9384-10MG |
| S Adenosyl-L -methionine | Sigma Aldrich | A7007-25MG |
| 5-Methyltetrahydrofolic acid | Sigma Aldrich | M0132-10MG |
| Betaine | Sigma Aldrich | B3501-100G |
| Choline | Sigma Aldrich | C7017-5G |
| Dimethyl glycine | Sigma Aldrich | D6382-5G |
| Homocysteine d4 | novachem (CIL) | DLM-8259 |
| Cystathionine d4 | novachem (CIL) | DLM-6108 |
| S-Adenosylhomocysteine d4 | Sapphire bioscience (Cayman) | 900037 |
| 5-Methyltetrahydrofolic acid | novachem (CIL) | CLM-7321-PK |
| Betaine d3 ( N-(Carboxymethyl)-N,N,N-trimethyl-d3-ammonium Chloride (N-methyl-d3) | SciVac PTY. Ltd. (CDN) | D-6303 |
| Dimehtyl glycine d3 | SciVac PTY. Ltd. (CDN) | D-7024 |
| 5-(Methyl-d3) tetrahydrofolic acid calcium salt | Sapphire Bioscience Pty Ltd | M330132 |
| Taurine d4 | SciVac PTY. Ltd. (CDN) | CDN Isotopes D-1971 |
| Betaine | Sigma Aldrich | B3501-100G |
| Choline | Sigma Aldrich | C7017-5G |
| Dimethyl glycine | Sigma Aldrich | D6382-5G |
| Taurine | Sigma Aldrich | T6025-10G |
| L- Alanine | Sigma Aldrich | A 7627 |
| L-Serine | Sigma Aldrich | S 4500 |
| Glycine | Sigma Aldrich | G 7126 |
| L-Glutamine | Sigma Aldrich | G 3126 |
| L-Proline | Sigma Aldrich | P 0380 |
| L-Lysine, HCl | Sigma Aldrich | L 5626 |
| L-Arginine, HCl | Sigma Aldrich | A 5131 |
| L-Valine | Sigma Aldrich | V 0500 |
| L-Threonine | Sigma Aldrich | T 8625 |
| Hydroxy-L-proline | Sigma Aldrich | H 6002 |
| L-Histidine, HCl | Sigma Aldrich | H 8125 |
| L-Asparagine | Sigma Aldrich | A 0884 |
| Taurine | Sigma Aldrich | T6025-10G |
| L-Glutamic acid | Sigma Aldrich | G1251 |
| Ornithine | Sigma Aldrich | O-2375 |
| L- Methionine | Sigma Aldrich | M 9625 |
| L-Tyrosine | Sigma Aldrich | T 3754 |
| L-Isoleucine | Sigma Aldrich | I 2752 |
| L-Leucine | Sigma Aldrich | L 8000 |
| L-Phenylalanine | Sigma Aldrich | P 2126 |
| L-Tryptophan | Sigma Aldrich | T 0254 |
| L-Aspartic acid | Sigma Aldrich | A 9256 |
| Amino adipic acid | Sigma Aldrich | A7275-100MG |
| Ethanolamine | Sigma Aldrich | 398136-25mL |
| Citrulline | Sigma Aldrich | C7629-1G |
| 1-methylhistidine | Sigma Aldrich | 67520-50MG |
| 3-methylhistidine | Sigma Aldrich | M9005-100MG |
| Carnitine | Sigma Aldrich | C0283-1g |

| **Table 2: Unlabelled 1-C metabolism mix** | | | |
| --- | --- | --- | --- |
| Compound | Volume (µL) to pipette | Final concentration (µM) | Stock concentration (µM) |
| Cysteine | 100 | 500 | 5000 |
| Homocysteine | 40 | 200 | 5000 |
| Cystathionine | 2 | 1 | 500 |
| S-Adenosylhomocysteine | 2 | 1 | 500 |
| S Adenosyl-L -methionine | 2 | 1 | 500 |
| 5-Methyltetrahydrofolic acid | 10 | 5 | 500 |
| Betaine | 100 | 500 | 5000 |
| Choline | 25 | 400 | 16000 |
| Dimethyl glycine | 80 | 400 | 5000 |
| Taurine | 200 | 1000 | 5000 |
| Carnitine | 200 | 1000 | 5000 |

| **Table 3: Unlabelled amino acid mix** | | | |
| --- | --- | --- | --- |
| Compound | Volume (µL) | Final concentration (µM) | Stock concentration (µM) |
| Alanine | 20 | 1000 | 45000 |
| Serine | 20 | 1000 | 45000 |
| Glycine | 20 | 1000 | 45000 |
| Glutamine | 20 | 1000 | 45000 |
| Proline | 20 | 1000 | 45000 |
| Lysine | 20 | 1000 | 45000 |
| Arginine | 20 | 1000 | 45000 |
| Valine | 20 | 1000 | 45000 |
| Threonine | 20 | 1000 | 45000 |
| OH-proline | 15 | 400 | 24000 |
| Histidine | 15 | 400 | 24000 |
| Asparagine | 15 | 400 | 24000 |
| Taurine | 15 | 400 | 24000 |
| Glutamic acid | 15 | 400 | 24000 |
| Ornithine | 15 | 400 | 24000 |
| Methionine | 15 | 400 | 24000 |
| Tyrosine | 15 | 400 | 24000 |
| Isoleucine | 15 | 400 | 24000 |
| Leucine | 22.5 | 600 | 24000 |
| Phenylalanine | 15 | 400 | 24000 |
| Tryptophan | 15 | 400 | 24000 |
| Ethanolamine | 30 | 400 | 12000 |
| 3-aminobutyric acid | 30 | 400 | 12000 |
| 2-aminoisobutyric acid | 30 | 400 | 12000 |
| Aminoadipic acid | 30 | 400 | 12000 |
| Aspartic acid | 30 | 400 | 12000 |
| Citrulline | 30 | 400 | 12000 |
| 1-methylhistidine | 30 | 400 | 12000 |
| 3-methylhistidine | 30 | 400 | 12000 |
